# X-Chromosomal STR-based analysis of genetic diversity, structure, and phylogeny in the Sri Lankan Vedda population

**DOI:** 10.64898/2026.10.26.761744

**Authors:** Shenali A. Ranasinghe, K. L. Nandika Perera, Gayani H. Galhena

**Author notes:** Corresponding authors: Prof. Gayani Harendra Galhena, Shenali Avishka Ranasinghe.

## Abstract

The Vedda people are regarded as the direct descendants of Sri Lanka’s Mesolithic hunter-gatherers and represent one of South Asia’s oldest surviving indigenous populations. X-chromosomal short tandem repeat (X-STR) analysis provides a powerful framework to examine their genetic distinctiveness, as the X chromosome exhibits sex-biased biparental inheritance and greater sensitivity to demographic and evolutionary forces compared to autosomal markers. However, no X-STR-based genetic investigations have yet been conducted on this population. In the present study, we successfully genotyped 50 unrelated Veddas individuals using a recently developed 16 X-STR multiplex PCR system optimized for Sri Lankan populations. Our analyses revealed a statistically significant genetic differentiation (2.43%) between the Veddas and the principal local ethnic groups—Sinhalese, Sri Lankan Tamils, Indian Tamils, and Moors— exceeding previous estimates based on autosomal STR markers. Comparative analysis with 32 global populations positioned the Veddas as a genetically distinct and isolated group, separated from clusters representing South Asians, Europeans, Middle Easterners, and other indigenous populations such as Australian Aboriginals, Bhil Indians, and Kedayans, emphasizing their unique genetic heritage.

Reduced allelic (115) and haplotype (62) counts, alongside significant linkage disequilibrium detected among eight pairs of X-STR loci, reflect the influence of genetic drift within this small, endogamous, and historically isolated population. These findings highlight the evolutionary and demographic distinctiveness of the Veddas and provide essential baseline data for future population genomic studies. Furthermore, the demonstrated high discriminatory power of the 16 X-STR loci supports their continued application in population genetics and anthropological research within this unique community.

## Introduction

Sri Lanka is an island nation-state of South Asia located near India. It remained a part of mainland Asia until a few thousand years back, facilitating the influx of anatomically modern humans as they moved out of Africa. Despite Sri Lanka’s geographical separation from Asia, it faced several colonial invasions and migrations, leading to its current ethnic diversity (1). The island is now inhabited by 22 million, of which 99.5% is accounted for by the four major ethnic groups of Sinhalese, Sri Lankan Tamils (SLT), Indian Tamils (IT) and Moors, while the Malays and Burghers together with the indigenous community account for less than 0.5% of the total population (2).

The indigenous community (aboriginals) of Sri Lanka are widely known as Veddas, which is derived from the Sanskrit word “Vyadha” depicting a hunter with a bow and arrow, thereby expressing their hunter-gatherer way of life. The origin of Veddas has been primarily debated due to the different narratives in ancient literature but is widely believed to be a continuation of the “Yakkas” (the indigenous tribe inhabiting the land during the earliest Indian invasions). Archaeological evidence from several inland cave sites demonstrated the presence of the earliest anatomically modern humans since 45,000 years ago, coinciding with the early human migrations (3,4). Due to morphological similarities such as thick skulls, heavy jaws, short necks, and robust bones between these early Mesolithic-hunter-gatherers and the Veddas, a long-preserved genetic continuum between the two populations has been suggested (5). This favours the possibility of Veddas representing a genetic continuum of these early modern humans and having their genetic similarities lying within the Australo-Melanesian aboriginals who are scattered in many parts of Asia (6).

Over time, with successive invasions and the expansion of new populations across Sri Lanka, the Veddas have become restricted to remote areas, eventually dwindling in number. They now live restricted to several regions in Sri Lanka such as “Dambana”, “Pollebadda” and “Rathugala”. As the lives of this ancient population of Veddas gradually integrate with the more dominant local ethnic groups, the danger of extinction looms over them, making it necessary to identify their genetic identity and affinities both in a local and global context. Several studies seeking the genetic structure of the Veddas have already been conducted utilising biochemical (7–10), autosomal (11) and mitochondrial markers (12–15). A common narrative of a majority of these studies has been of the “Vedda” community being genetically differentiated from both local and global ethnic populations due to their long isolation from outside populations, while sharing slight genetic similarities with the Sinhalese, SLT and having West-Eurasian, Indian ancestries.

However, none of these previous studies had utilised markers on the X-chromosome, which display characteristics of both uniparental and biparental markers by undergoing recombination within a smaller effective population size (16). This allows less space for mutations but more prone to evolutionary forces such as genetic drift, selection and gene flow compared to autosomes. Particularly, short tandem repeat (STR) markers display greater polymorphism and are capable of detecting genetic drift within small, isolated populations (17). Therefore, X chromosomal STRs (X-STRs) appear as a large asset in the study of molecular anthropology, even among indigenous communities, to reveal their sex-biased demographic history (18). In addition to evolutionary studies, X-STRs are also useful in forensic genetics owing to their high discriminatory power and the presence of tightly linked STRs forming highly informative haplotypes. They, therefore have the potential to be successfully used in the resolution of complex kinship cases involving at least one female (19). The scope of X-STRs in population and forensic genetics has already been investigated in several global populations, including indigenous communities but is yet to be investigated for the Veddas.

An inhouse 16 X-STR multiplex PCR system has already been developed and validated in Sri Lanka based on the main Sri Lankan ethnicities and is being used in routine forensic genetic applications (20,21). The X-STRs utilised consist of 13 X-STR loci that occur as four clusters (Cluster I; DXS10148-DXS10135-DXS8378 (Xp22), Cluster II; DXS7132-DXS10079-DXS10074-DXS10075 (Xq12), Cluster III: DXS6801-DXS6809-DXS6789 (Xq21), Cluster IV; DXS7424-DXS101-DXS7133 (Xq22)) and three single independent loci, one of which lies on the p arm (DXS9902 at Xp22), with the other two on the q arm (HPRTB at Xq26 and DXS7423 at Xq28). These STRs span almost the entire length of the X−chromosome from 9.198 Mb to 149.46 Mb, allowing better coverage of the chromosome. The 16 X−STR multiplex system contains eight loci in common with the popularly used Argus X−12 kit, which allows allele frequency comparisons with global population genetic data.

The present study analyses the Sri Lankan aboriginals utilising these 16 X-STR loci and reports for the first time, their X-STR-based population genetic data. Furthermore, it compares their genetic positioning relative to the four main Sri Lankan ethnicities (Sinhalese, SLT, IT and Moors) and 32 global populations including several indigenous communities (Australian aboriginals, Kedayans, Bhil Indians, Gran Chaco Amerindians), based on pairwise genetic distance calculations. Results also include those of a comprehensive analysis of linkage disequilibrium (LD) conducted for the selected X-STR markers on the aboriginal community and report forensic efficiency of the 16 X-STR multiplex system while constructing a separate allele and haplotype frequency database for the “Vedda” people of “Dambana”, Sri Lanka for purposes of forensic and kinship analyses.

## Materials and Methods

### Sample preparation and DNA extraction

Ethical clearance for sample collection was obtained from the Ethics Review Committee of the Institute of Biology, Sri Lanka (ERC-IOBSL-269-08-2022) and the research was conducted in line with the principles of the Declaration of Helsinki, where informed consent was obtained from participants of the study. Finger-pricked blood samples were collected from 50 unrelated Veddas (50% males) in “Dambana”, Sri Lanka. The sample size represents 0.5% of the total Vedda population in the area. Chelex-100 method was used in genomic DNA extraction and amplified using the single tube 16 X-STR multiplex PCR system (21). Resolution of the amplified product was carried out by capillary electrophoresis using the ABI 3500 Genetic Analyzer (Thermo Fischer Scientific, USA). Genotyped data were collected using 3500 Series Data Collection Software 3 (Thermo Fischer Scientific, USA) and allele designation was performed by the GeneMapper IDX software (Thermo Fischer Scientific, USA).

### Population genetic data analysis

Of the 50 samples, two male samples had almost similar profiles and therefore one was removed from the database and analysis due to the possibility of relatedness. Allele and haplotype frequency calculations, exact test of differentiation between male and female allele frequencies, compliance of female genotype frequencies with the Hardy Weinberg Equilibrium, and the test of pairwise linkage-disequilibrium (LD) within and between clusters of linked loci of male samples were analyzed using ARLEQUIN 3.5.2. Genetic structuring of Veddas against the four main ethnic groups comprising of migrants that settled in Sri Lanka after 543 B.C. (i.e., Sinhalese, SLT, IT and Moors) was analyzed using previously published data (20) through locus-by-locus Analysis-of-Molecular-Variance (AMOVA) on ARLEQUIN 3.5.2. Population-specific inbreeding co-efficient values (Fis) were calculated for the Veddas and four local ethnicities at 10,000 permutations. Pairwise genetic distances (Fst) were also calculated for all pairs of populations based on Nei’s average number of pairwise differences within and between populations, where the null distribution of pairwise Fst was obtained by permuting haplotypes between populations 1000 times at 0.05 significance. Genetic distances were visualized via a dendrogram constructed using POPGENE v.1.32 through the Unweighted-Pair-Group-Method-with-Arithmetic-mean (UPGMA). Fst analysis was conducted between Veddas and 27 aboriginal and other global populations using 8 common X-STR markers based on published data and visually represented using a multidimensional-scaling (MDS) plot drawn using IBM SPSS Statistics 26. Genetic admixture between the four Sri Lankan ethnicities, the Veddas and the Australian aboriginals was inferred using STRUCTURE v. 2.3.4, where cluster numbers (K) were tested from 1 to 6 with 5 runs per K, a burnin length of 100,000 followed by 500,000 Markov-Chain-Monte-Carlo (MCMC) iterations, using the ADMIXTURE model with LOCPRIOR on and correlated allele frequencies. Optimal K was obtained upon calculation of Delta K.

### Forensic data analysis

Parameters of forensic interest, namely polymorphism-information-content (PIC), expected heterozygosity (Het), mean-exclusion-chance-in-deficiency (MEC_Kru⃛ger_), mean-exclusion-chance-in-Duos (MEC_Desmarais duo_), mean-exclusion-chance-in-trios (MEC_Desmarais trio_), power-of-discrimination-for-females (PD_f_) and power-of-discrimination-for-males (PD_m_) were calculated for all 16 X−STR loci as well as the haplotypes as defined by the four clusters using MATLAB software (version R2021a) and chromosome X web (http://www.chrx-str.org). Forensic parameters were also calculated for loci under linkage disequilibrium based on their haplotype frequencies using the same platforms.

## Results

### STR genotyping

Complete DNA profiles of all Veddas were generated with no allele dropouts, giving a total of 115 alleles from the 16 X−STR loci (Supplementary table 1). This, however, was less compared to the other local ethnicities of Sinhalese (184), SLT (164), IT (154) and Moors (150) (20).

All loci of female samples showed no deviations from the Hardy-Weinberg equilibrium (HWE) expectations after correcting for multiple comparisons (Bonferroni-corrected-P= 0.0031), indicating random assortment of alleles and the lack of genotyping errors, confirming their selective neutrality (Supplementary table 2). Heterozygosity at all loci exceeded 0.5, informing its applicability in studying population structure owing to the high discriminatory power (17). However, the average observed heterozygosity of the Veddas (0.7350) was slightly lower than that reported previously (20) among the Moors (0.7540), Sinhalese (0.7537) and IT (0.7471).

The exact test of population differentiation comparing the allele distribution between male and female aboriginals did not indicate any significant differences (P=1.00), and therefore, allele frequencies of both sexes were combined for further analysis.

### Population differentiation among Sri Lankan ethnicities

Genetic structuring of the Veddas among the four main Sri Lankan ethnicities of Sinhalese, SLT, IT and Moors was assessed using locus-by-locus Analysis-of-Molecular-Variance-Framework (AMOVA), where differences in allelic content of all 16 X-STR loci were taken into consideration (Supplementary Table 3). Here, the four main ethnicities were considered as one group comprised of migrants who were historically known to have settled in Sri Lanka much later in comparison to the Veddas. The Sinhalese, descendants of the Indo-Aryans who migrated from North India in 543 B.C. (22), the SLT of Dravidian origin who settled during the South Indian expansions to Sri Lanka (6), Moors who were descendants of either Arabian traders or South Indian Muslims (23) and IT brought in by European colonists as estate labourers from South India (1) were grouped together. Interestingly, a statistically significant differentiation of 2.43% was identified between the two groups, demonstrating a clear population subdivision between the Veddas and the four main local ethnicities. A ten-fold lower but significant variation of 0.20% existed among the four ethnicities within the group, while 1.74% of the variance in the samples was attributable to that within different populations. Pairwise population comparisons via Fst analysis were also conducted for the five populations to better understand the above population structure (Table 1). Accordingly, the Veddas showed a similar statistically significant genetic subdivision from all the other ethnicities. Even though the only population pair with no significant genetic differentiation were the Sinhalese and Moors, differentiation between the aboriginals with each of the four ethnicities stood ten times greater than any of the other pairs. The UPGMA phylogram (Figure 1) visualised the genetically distant relationship between the Veddas and the four main ethnicities, all of which had close genetic affinities based on Nei’s genetic distance calculations (Supplementary table 4). The Veddas appeared as an outgroup while the Sinhalese and Moors showed close genetic associations, followed by the SLT and IT that occurred towards the periphery of the internal cluster.

**Figure 1:**
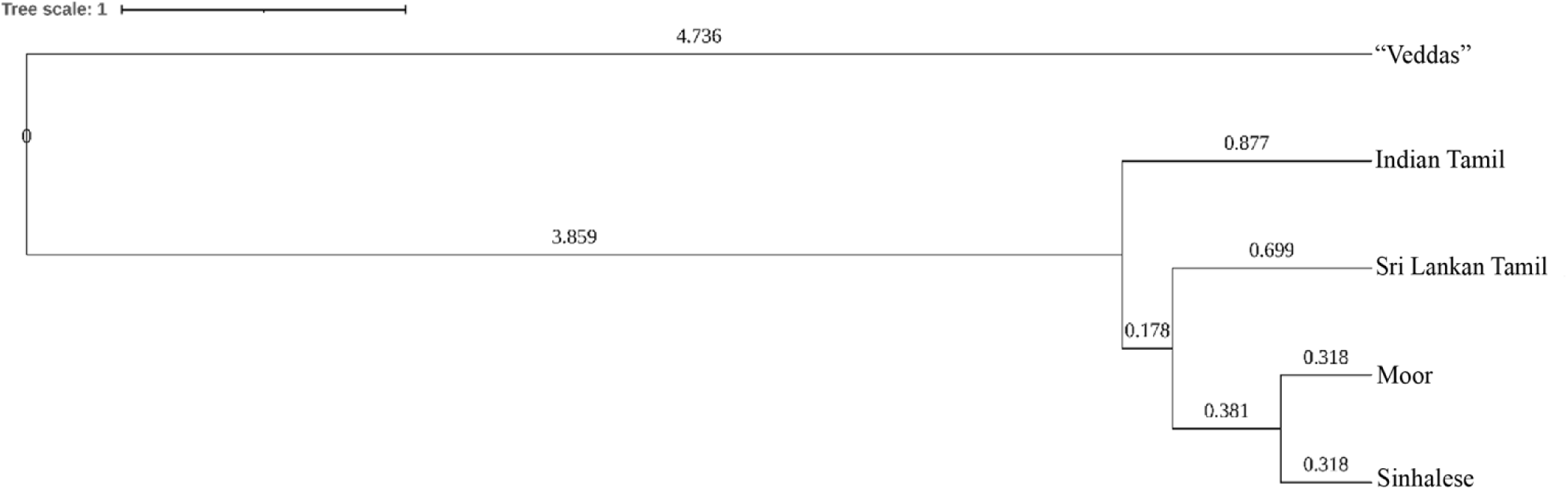
UPGMA phylogram of Veddas and the four Sri Lankan ethnicities based on 16 X-STR data. Branch lengths are in the same units as the evolutionary distances used in inferring the phylogenetic tree.

**Table 1:**
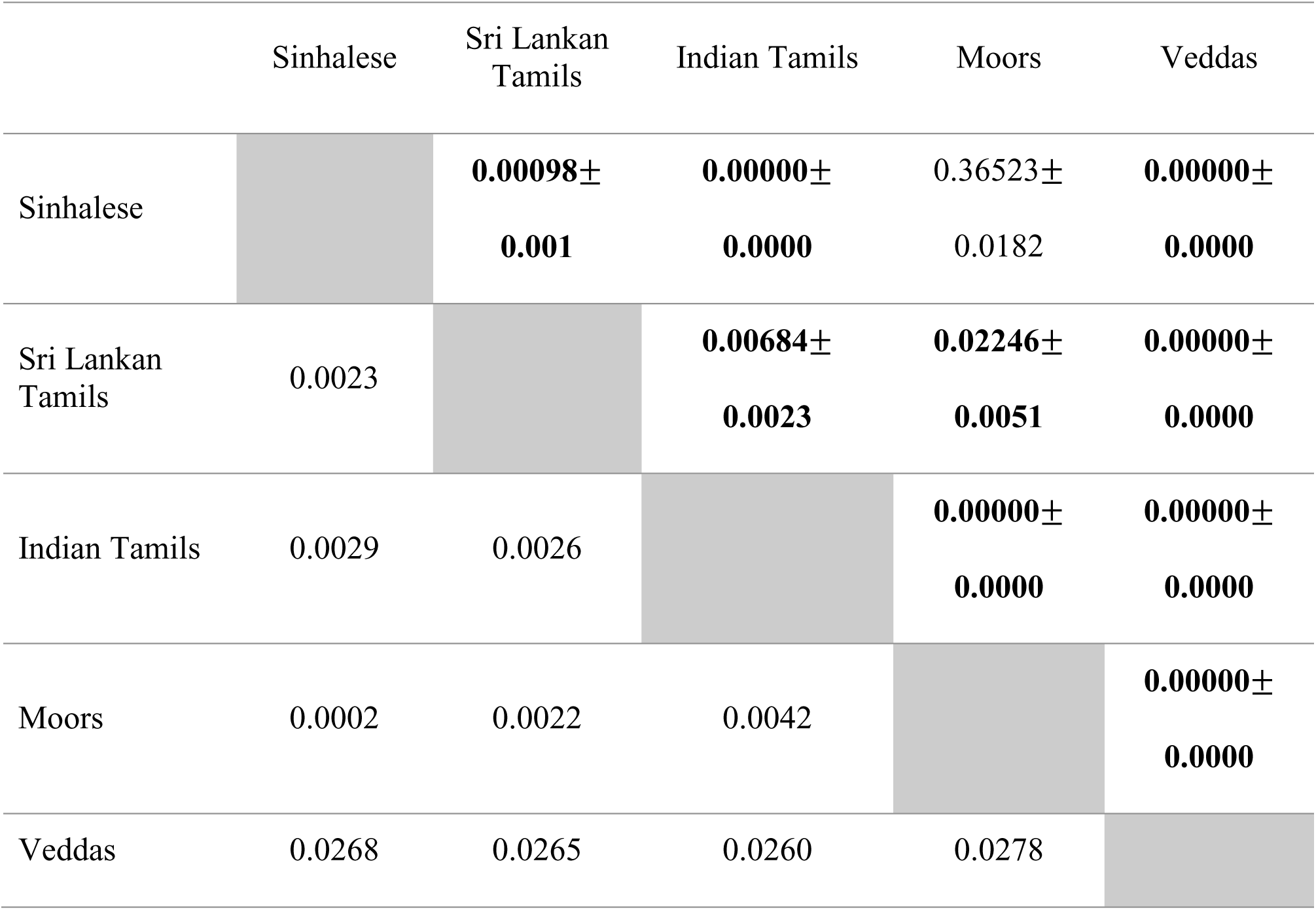
Pairwise population differentiation (Fst) and corresponding P-values for the Veddas and the four main Sri Lankan ethnicities. Below diagonal: pairwise Fst values, above diagonal: P-values with significant P-values in bold.

|  | Sinhalese | Sri Lankan Tamils | Indian Tamils | Moors | Veddas |
|---|---|---|---|---|---|
| Sinhalese |  | **0.00098± 0.001** | **0.00000± 0.0000** | 0.36523± 0.0182 | **0.00000± 0.0000** |
| Sri Lankan Tamils | 0.0023 |  | **0.00684± 0.0023** | **0.02246± 0.0051** | **0.00000± 0.0000** |
| Indian Tamils | 0.0029 | 0.0026 |  | **0.00000± 0.0000** | **0.00000± 0.0000** |
| Moors | 0.0002 | 0.0022 | 0.0042 |  | **0.00000± 0.0000** |
| Veddas | 0.0268 | 0.0265 | 0.0260 | 0.0278 |  |

Inbreeding coefficient (Supplementary table 5) of the Veddas was a statistically non-significant negative value (Fis= –0.0047; P>0.05), depicting the possible absence of inbreeding. Though all other ethnicities showed positive inbreeding, only the SLT exhibited statistically significant Fis (Fis= 0.0430; P<0.01).

### Genetic positioning of the Veddas among the global populations

Pairwise Fst values generated for Veddhas of the present study and 27 other populations (Sinhalese, SLT, IT and Moors (20), Australian aboriginals (24), Jat Sikhs (25), Kedayans of Borneo (26), Egypt (27), Saudi Arabia (28), Greenland (29), Guinea Bissau (30), Algeria (31), Mexico (32), and 14 additional populations cited in Perera et al., (20) for whom data were available are presented in Supplementary table 6. When the results are visualised in a 2D-MDS plot (Figure 2), the Veddas occupied an isolated position with no close link to any of the local and global populations. They were positioned further away from the large central cluster formed by the Sri Lankan, South Asian, European, Middle-Eastern populations, and Australian aboriginal population. Southeast Asians, East Asians, Africans, Mexicans and Greenlanders did not occur within this cluster and showed no close relationship with the Veddas either.

**Figure 2:**
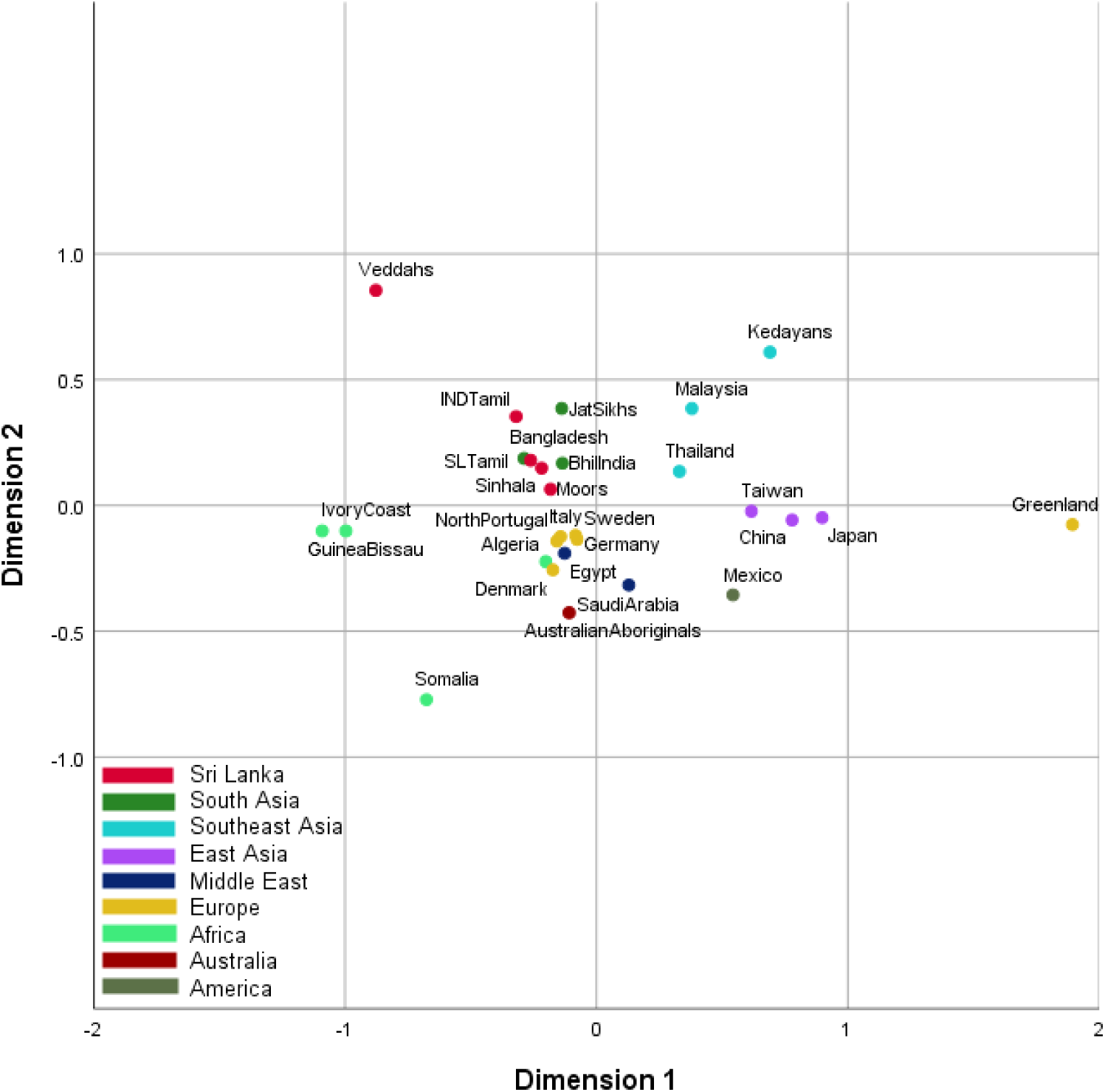
Two-dimensional MDS plot illustrating the relationship between Veddas and 27 local and global populations using pairwise Fst values averaged over eight common X-STR loci.

Since STRUCTURE calculates genetic admixture based on the assumptions of loci being at HWE and linkage equilibrium (33), the analysis was conducted using only 10 X−STRs with no LD (DXS10148, DXS8378, DXS9902, DXS7132, DXS7132, DXS6801, DXS6789, DXS7424, DXS7133, HPRTB and DXS7423), which indicated the most likely presence of three genetic clusters (K=3) according to the Evanno method (34). The four local ethnicities (blue), Veddas (green) and Australian aboriginals (red) formed 3 clusters with clear separation (Figure 3). The Veddas showed a unique genetic composition with slight admixture from the local ethnicities and Australian aborigines. This indicated that the Sinhalese, SLT, IT, and Moor populations shared genetic similarities and had a distinct genetic profile compared to the Veddas, which aligned with the phylogenetic tree from Figure 1. The noticeable presence of Vedda ancestry among a few SLT and IT individuals may reflect relatively recent isolated cases of admixing. The Sinhalese and more prominently the Moors share genetic similarities with the Australian aboriginals than the Veddas, possibly indicating their West European admixture. However, this accounted for only a small proportion of their tested genetic composition.

**Figure 3:**
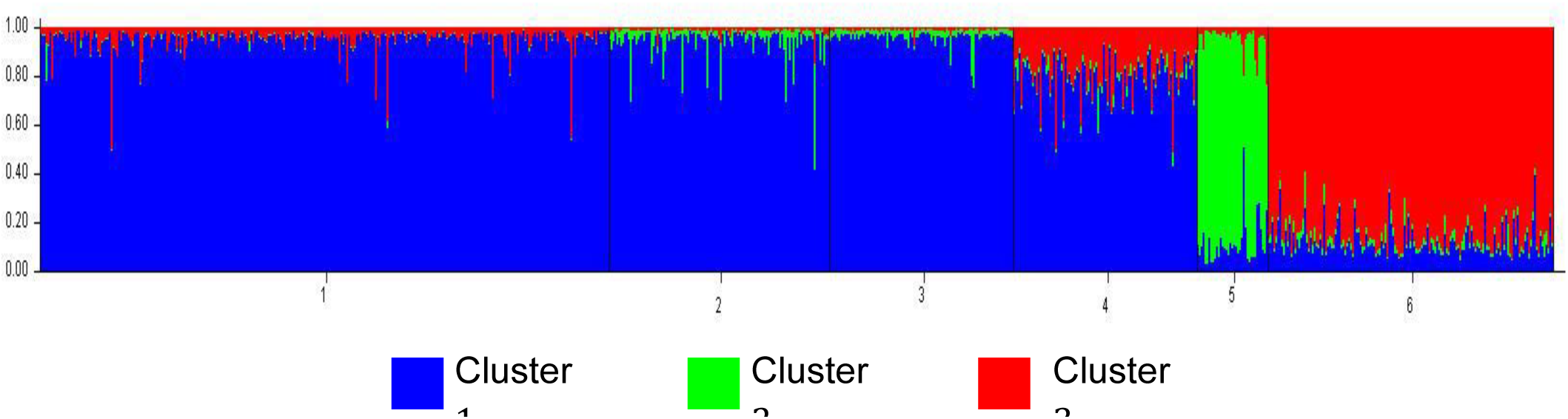
Population structure of the four Sri Lankan ethnicities, Veddas and Australian aboriginals at K= 3 as inferred by the STRUCTURE v. 2.3.4 program after 100,000 burnin and 500,000 MCMC replications. Y axis corresponds to the inferred ancestry from the four clusters. Each individual is represented by the vertical axes and the six populations are separated by the black lines. 1 −Sinhalese, 2 −Sri Lankan Tamils, 3 −Indian Tamils, 4 −Moors, 5 −Veddas, 6 − Australian aboriginals.

### Linkage disequilibrium and haplotype analysis

LD is a population-specific phenomenon that is affected by a variety of evolutionary forces, such as founder effects, population bottlenecks, random genetic drift, and inbreeding, which are helpful in revealing a population’s demographic history (35). Hence, LD of the Veddas was calculated between each pair of markers in the four separate clusters (Table 2).

**Table 2:** Pairwise LD results of the Veddas. Statistically significant P values after correcting for multiple comparisons (Bonferroni corrected P= 0.0167, P= 0.0125, P= 0.0167, P= 0.0167, and P= 0.0083 for each cluster/ cluster pair in the same order as given in the table) given in bold font.

| Loci pair | P value |
|---|---|
| **Cluster I** |  |
| DXS10148, DXS10135 | **0.0000** |
| DXS10148, DXS8378 | 0.4218 |
| DXS10135, DXS8378 | 0.0401 |
| **Cluster II** |  |
| DXS7132, DXS10079 | 0.0556 |
| DXS7132, DXS10074 | 0.2349 |
| DXS10079, DXS10074 | **0.0021** |
| DXS7132, DXS10075 | **0.0063** |
| DXS10079, DXS10075 | **0.0000** |
| DXS10074, DXS10075 | **0.0014** |
| **Cluster III** |  |
| DXS6801, DXS6809 | 0.3563 |
| DXS6801, DXS6789 | 0.7148 |
| DXS6809, DXS6789 | **0.0002** |
| **Cluster IV** |  |
| DXS7424, DXS101 | **0.0091** |
| DXS7424, DXS7133 | 0.1006 |
| DXS101, DXS7133 | **0.0018** |
| **Clusters III and IV** |  |
| DXS6801, DXS7424 | 0.4632 |
| DXS6809, DXS7424 | 0.1623 |
| DXS6789, DXS7424 | 0.0283 |
| DXS6801, DXS101 | 0.1433 |
| DXS6809, DXS101 | 0.1220 |
| DXS6789, DXS101 | 0.2578 |
| DXS6801, DXS7133 | 0.3061 |
| DXS6809, DXS7133 | 0.8518 |
| DXS6789, DXS7133 | 0.3644 |

Significant LD was detected between eight pairs of X−STR markers among Veddas, and this was more profound than reported in any of the four local ethnicities (20). Within cluster I, LD with a very high significance was detected only between DXS10148-DXS10135, which are 1 kb apart.

Cluster II had the highest occurrence of LD among all four clusters, between DXS10079-DXS10075, DXS10079-DXS10074, DXS7132-DXS10075 and DXS10074-DXS10075, some of which were not even adjacent loci. Cluster III only had a single pair of adjacent markers DXS6809-DXS6789 with significant LD and cluster IV had two pairs of markers DXS7424-DXS101 and DXS101-DXS7133 both of which were placed directly adjacent to one another. Moreover, pairwise LD between clusters III and IV was also analysed due to their close physical proximity (6.78 cM), but no significant LD was detected in any of the marker pairs.

### Haplotype frequencies

Haplotype frequency calculations of the Veddas were performed using only the male samples since the hemizygous nature of their X−chromosome allow direct haplotype estimations (Supplementary table 7). Interestingly, each sampled male produced a unique haplotype with respect to all markers, thus indicating the high discriminatory power of the 16 X−STR assay system in characterising the individuals of the Vedda population. Moreover, cluster I was found to be the most informative in comparison to all other clusters with a haplotype diversity of 0.9818, due to the presence of DXS10148 and DXS10135 both of which are highly polymorphic loci. This was also found to be consistent with the other four local ethnicities, with the Veddas having a lower haplotype diversity in comparison.

However, due to the occurrence of LD within the four clusters, haplotype frequencies were also calculated for the loci as clusters (Supplementary tables 8, 9), giving a total of 62 haplotypes for the Veddas. Haplotype diversity among the aboriginals was highest in cluster I (0.9818) and lowest in cluster IV (0.8983), a pattern similar to that observed in the other four ethnicities. A few of the markers considered under clusters II and IV in this study, showed variations from previous publications. STRs of cluster II were considered in two different combinations as; DXS7132-DXS10079-DXS10074 (Argues X-12 kit) and DXS10079-DXS10074-DXS10075 (36) and STRs of cluster IV considered to comprise of only two markers; DXS7474-DXS101 disregarding DXS7133 (37). Therefore, haplotype frequencies were calculated for these cluster combinations as well (Supplementary table 10).

### Forensic efficiency parameters

Owing to the unique genetic composition thus far observed within the Veddas, already available population frequency databases would not be appropriate for kinship or forensic casework concerning them. More exhaustive sampling to include Veddas from other sub-groups across different geographic areas within Sri Lanka would be essential in constructing a representative allele frequency database, which can be used to accurately calculate expected genotype frequencies, match probabilities and likelihood ratios.

Forensic efficiency parameter calculations help verify the applicability of the tested X-STR loci for the Veddas in forensic and kinship applications. Since the haplotypes that occur in loci with LD tend to behave as alleles (i.e., single locus), forensic efficiency parameters were calculated based on the haplotype frequencies instead of individual allele frequencies (Table 3). Accordingly, PIC, Het, PD_f_ and PD_m_values for the haplotype DXS10148-DXS10135 exceeded 0.9 while MEC values ranged above 0.8. In the other three haplotypes defined by LD, the PIC, Het, PD_f_ and PD_m_ values surpassed 0.8 and showed values ranging from 0.6-0.9 for the MEC parameters. Thus, the PIC, Het, PD_f_ and PD_m_ values confirmed the 16 X-STR markers to be informative and sufficiently discriminative for the Veddas. MEC_Desmarais trio_ and MEC_Desmarais duo_ also indicated the suitability of the X−STR panel in resolving kinship in trios with a female offspring and father-daughter or mother-son duos, respectively. Combined forensic efficiency parameters showed that the combined power of discrimination for females (CPD_f_) and males (CPD_m_), combined MEC indices for deficiency (CMEC_Krüger_), normal trio (CMEC_Desmarais−trio_) and duo cases (CMEC_Desmarais−duo_) all came close to one. These results therefore confirmed the suitability of the 16 X−STR assay in kinship and forensic investigations of the Veddas.

**Table 3:**
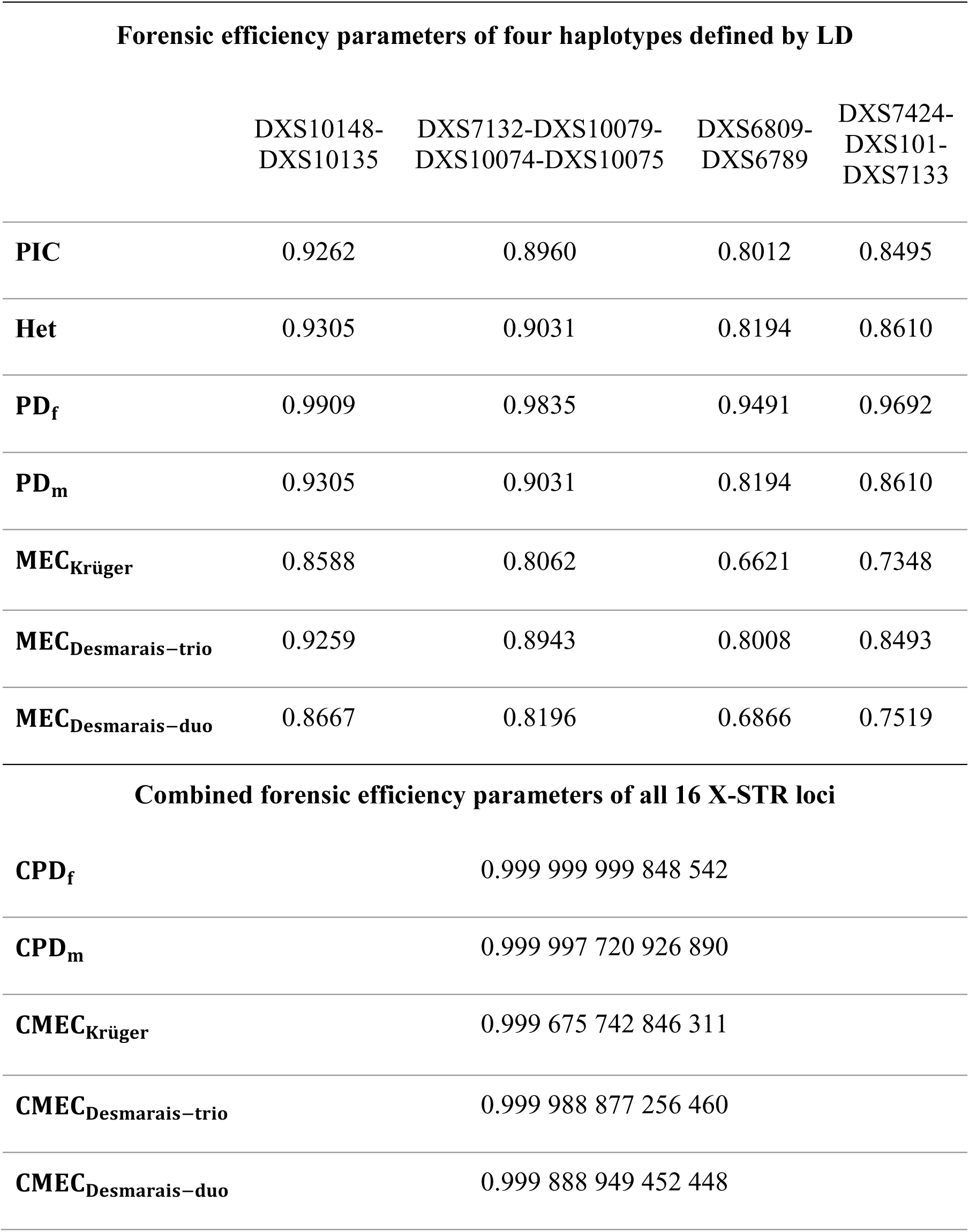
Forensic efficiency parameters of four haplotypes defined by LD and combined forensic efficiency parameters for the 16 X-STR loci of the Veddas. PIC polymorphism information content, Het expected heterozygosity, MEC_Kru⃛ger_mean exclusion chance in deficiency, MEC_Desmarais duo_ mean exclusion chance in Duos, MEC_Desmarais trio_ mean exclusion chance in trios, PD_f_ power of discrimination for females and PD_m_ power of discrimination for males.

| **Forensic efficiency parameters of four haplotypes defined by LD** |  |  |  |  |
|  | DXS10148-DXS10135 | DXS7132-DXS10079-DXS10074-DXS10075 | DXS6809-DXS6789 | DXS7424-DXS101-DXS7133 |
|---|---|---|---|---|
| **PIC** | 0.9262 | 0.8960 | 0.8012 | 0.8495 |
| **Het** | 0.9305 | 0.9031 | 0.8194 | 0.8610 |
| **PD<sub>f</sub>** | 0.9909 | 0.9835 | 0.9491 | 0.9692 |
| **PD<sub>m</sub>** | 0.9305 | 0.9031 | 0.8194 | 0.8610 |
| **MEC<sub>Krüger</sub>** | 0.8588 | 0.8062 | 0.6621 | 0.7348 |
| **MEC<sub>Desmarais−trio</sub>** | 0.9259 | 0.8943 | 0.8008 | 0.8493 |
| **MEC<sub>Desmarais−duo</sub>** | 0.8667 | 0.8196 | 0.6866 | 0.7519 |
| **Combined forensic efficiency parameters of all 16 X-STR loci** |  |  |  |  |
| **CPD<sub>f</sub>** | 0.999 999 999 848 542 |  |  |  |
| **CPD<sub>m</sub>** | 0.999 997 720 926 890 |  |  |  |
| **CMEC<sub>Krüger</sub>** | 0.999 675 742 846 311 |  |  |  |
| **CMEC<sub>Desmarais−trio</sub>** | 0.999 988 877 256 460 |  |  |  |
| **CMEC<sub>Desmarais−duo</sub>** | 0.999 888 949 452 448 |  |  |  |

## Discussion

Given the sex-biased inheritance and smaller effective population size of the X-chromosome, together with the maternal inheritance of mtDNA, previous findings based on mtDNA (12–15) align with the current observations as they also indicate the Veddas to be genetically differentiated from the rest of the populations who have settled in the island much later based on the recorded history. This supports the hypothesis of the Veddas being descendants of the early inhabitants of the island. The only other STR analysis conducted on the Veddas was based on autosomes and suggests a 2.1% genetic differentiation between them and the local ethnicities (11), confirming the slightly higher sensitivity of the 16 X−STR multiplex system in detecting population sub-structure. This may have been due to the greater capability of haploid loci (the X chromosome acts as a haploid locus in males) with low effective population sizes in capturing population divergence much better compared to the biparental autosomal loci with relatively larger effective population size (38). However, given that the above-mentioned studies have included samples from various Vedda sub-groups corresponding to different geographical localities within the island, direct comparisons cannot be made. This also indicates the scope to which the current analysis can be expanded to explore the intra-population structure of different Vedda populations using X-STRs.

Being a small population themselves with the gradual decline in their population size, the Veddas would have been affected by population bottlenecks and substantial genetic drift similar to many other tribal populations (39), leading to their lower genetic diversity. Furthermore, while the four main ethnicities underwent continued gene flow between each other as well as with the global populations, the Veddas adhered to a largely endogamous lifestyle, causing genetic isolation. Genetic drift and the gradual accumulation of mutations would have resulted in their characteristic allele frequencies and unique genetic makeup, making the Veddas an extreme outlier in the Sri Lankan ethnic phylogeny. However, the observed possible absence of inbreeding within the population upon Fis analysis could suggest that the population has remained sufficiently large and stable over time to avoid inbreeding.

During the out-of-Africa migration of early anatomically modern humans, they are known to have taken a coastal migratory route from Africa to arrive in the Indian subcontinent and subsequently Sri Lanka (40). As inferred from previous studies, if the Veddas are direct descendants of these early modern humans, the genetic continuum could be detectable upon comparison with other indigenous communities of the world. However, the MDS plot displayed otherwise, with the Veddas having no genetic affinities to either one of the global indigenous communities included in the comparisons namely, the Bhil tribe of India, Australian aborigines, and the Kedayans. This could have been caused by the extensive intermixing of the other aborigines compared to the Veddas, as well as the genetic drift and bottleneck effects over time. For instance, the Bhil Indians clustered closer together with the local Sri Lankan and South Asian populations instead of the Veddas. Previous studies analysing Y chromosomal and mitochondrial haplogroups show evidence of interbreeding between Indian aboriginal groups with expanding populations of the Fertile Crescent, creating the Aryan and Dravidian populace in India, who separate themselves from the aboriginals, eventually contributing largely to the later colonisations of Sri Lanka (41,42). Hence, the possible substantial admixture that must have occurred between the aboriginals of India and the “modernised” Indians must have caused the relative spatial positioning of the Bhil Indians away from the genetically isolated Veddas and closer to the South Asian populace. The settlement of Australian aborigines via Sahul took place approximately 50,000 years ago following their split from the African gene pool according to the single dispersal model (24) giving sufficient time for genetic divergence from the Veddas. During this period, intermixing of the Australo-Papuan ancestors (ancestors of Australian and New Guinean aboriginals) has occurred with the archaic hominin, Denisovans followed by extensive West European admixture of the Australian aboriginals during the British colonisation (43). Even the spatial positioning of the Australian aboriginals in the MDS plot depicts this close association with the Europeans and therefore would explain their divergence from the Veddas, who have not essentially undergone any known admixture as such. The Kedayans, an indigenous group in Borneo, depicted a similarly distant relationship with the Veddas. Previous X-STR studies conducted on this community (26) place their genetic affinities among the other Austronesian populations within Southeast Asia, while being dissociated from the European and African populations, similar to the current study.

However, it is notable that the Bhil Indians, Australian aboriginals and Kedayans significantly differentiated from the Veddas at only two, two and one loci respectively out of eight X−STR loci. Therefore, the relative positioning of the Veddas on the MDS plot was the result of both the absence of genetic affiliations between Veddas and other populations as well as the genetic proximity they shared with the other populations over the Veddas. Upon increasing the number of X−STR loci, however, more robust inferences regarding population differentiation could be made.

The findings from the Fst analysis and MDS plot, further aligned well with previous genetic studies comparing the Veddas and global populations. Kshatriya (8) studied genetic affiliations between the Veddas and populations spanning India, Middle East and Europe using 40 alleles controlled by 13 codominant loci, where in the dendrogram the Veddas were found to form an outgroup while the Sinhalese, Sri Lankan and IT, Gujaratis, Punjabis, Bengalis, Middle Eastern and European populations formed the inner branches in the given order. Hence, despite the use of first-generation markers such as blood markers, the clustering results agreed with those of the current study. However, no significant genetic differentiation between any of the populations was discovered, which could have been due to the use of conserved markers with low discriminatory power compared to the STR markers used in the current study. On the other hand, studies conducted using mtDNA hypervariable segments revealed the presence of West Eurasian haplogroups within the “Vedda” populations (12,14,15). According to this previous research, the presence of South Asian haplogroups within the Vedda populations has been detected, with remnants of N macro-haplogroups all of which arrived via the Southern migration route into Sri Lanka. Other STR-based studies also suggest slight genetic affinity between the Veddas and Sinhalese (11). Our Fst analysis also agrees with previous research and indicates a genetic relationship between the Veddas and the Sri Lankan and South Asian populations while strongly differentiating them from the East Asian, African and American populations.

Also notable was the presence of local aboriginal admixture within the SLT and IT in comparison to the others (i.e., Sinhalese and Moors). These Dravidian speakers of Sri Lanka are said to share closer genetic ties with the descendants of Ancestral Ancient South Indian (AASI) populations who represent early hunter-gatherers while the Sinhalese have roots within the Ancestral North Indians (ANI) who have expanded from Eurasian populations (1). Upon study of non-metric dental traits of Veddas, similarities with the Mesolithic South Asians have been discovered (44). Therefore, this could be the reason for the comparatively greater admixture between the Veddas and Tamils of Sri Lanka. However, some claim the possibility of the earliest AASI groups occupying Sri Lanka having been replaced by the ANI populations (1) due to the Eurasian links of the Veddas identified by previous research (14,15). This has been linked with the close resemblance of cranial shape that Veddas share with Dynastic Egyptians (45), North Indians and Middle Easterners (46) in proposing the ancestry of Veddas being rather more recent and connected with the ANI (1). However, no such higher admixture has been indicated between the Sinhalese and Veddas in comparison to the Dravidians and Veddas through the STRUCTURE plot. Therefore, in the presence of various dental, cranial and genetic evidence pointing in varying directions regarding their ancestry, whole genomic comparisons of Veddas, South Indians, North Indians and remains of the mesolithic men of India and Sri Lanka would help gather thorough evidence in narrowing down on one hypothesis regarding their origin (22).

With the Veddas being a small, reproductively isolated population, they can be subjected to the effects of genetic drift, which would then result in the loss of haplotypes, leading to increased LD (35). The original inhabitants of the Americas, the Amerindians, have also presented characters similar to the Veddas, such as high interpopulation divergence, low genetic diversity, and prevalence of LD (47), which would further support the above hypothesis. When comparing LD detected within the Veddas to that of other global aboriginal populations, such as the Australian aboriginals (24), Kedayans (26), Indian Bhils (48) and Native Central Americans (49), all have displayed a comparatively low occurrence of LD. Sparse LD has been documented in clusters I and II, with none in either clusters III or IV in all of the above populations. However, since the loci composition of the ARGUS X-12 kit used in these studies was different from that of the current study, direct comparisons are difficult. In adequately detecting LD, sample sizes exceeding 200 are given as a requirement since smaller sample sizes tend to artificially increase the presence of LD. Additionally, it is important to consider markers in linkage together instead of independently, when being applied in population genetic studies and forensic investigations (30). This would apply to the eight pairs of X−STR markers showing LD in the Veddas, however should be confirmed with a more representative sample.

Moreover, the total number of haplotypes was also lower in the Veddas in comparison to the other ethnicities and could have been a sign of the population bottleneck the Veddas were faced with (50). Alternatively, it might have been caused solely by the genetic drift since small, isolated populations tend to be subjected to significant effects of random genetic drift, leading to haplotype loss (35).

## Conclusion

This study represents the first investigation of X-STR variation in the largest extant aboriginal population of Sri Lanka, the Veddas of Dambana, with the sampled individuals accounting for approximately 0.5% of the Dambana community’s estimated population. Utilizing a 16-locus X-STR multiplex assay optimized for local relevance, we identified significant genetic differentiation between the Veddas and Sri Lanka’s major ethnic groups: Sinhalese, SLT, IT, and Moors. These findings highlight the utility of X-chromosomal markers in resolving fine-scale population structure, particularly within historically isolated and endogamous groups.

MDS analysis of pairwise genetic distances across 27 global populations revealed a highly distinct genetic profile for the Veddas, indicative of long-term genetic isolation. STRUCTURE analysis further supported this interpretation, revealing minimal admixture and reinforcing the population’s relative insulation from external gene flow. The observed patterns of reduced allelic and haplotype diversity, alongside widespread linkage disequilibrium, are consistent with the effects of historical bottlenecks, genetic drift, and prolonged small effective population sizes.

Although the present study does not conclusively resolve the ancestral origins of the Veddas, the findings provide strong evidence of their unique genetic legacy and underline their relevance in reconstructing early human population history in South Asia. The distinctiveness of their genomic profile raises the possibility of deep ancestral continuity, potentially extending to early modern human dispersals out of Africa.

While the primary focus of this research lies in elucidating evolutionary and demographic histories, the performance of the X-STR multiplex system also supports its potential value in kinship inference and population-specific forensic applications. Future studies incorporating broader geographic sampling, extensive genome-wide data, and ancient DNA will be critical in refining our understanding of the Vedda population’s demographic trajectory and its place within the broader framework of human genetic diversity.

Overall, our findings contribute an important layer to the growing understanding of human genetic diversity in South Asia and emphasize the value of preserving the genomic heritage of endangered indigenous populations.

## Acknowledgements

We are grateful to the Vedda Chief of Dambana, Uruwarige Wannila Aththo, for his invaluable support and consent, which enabled us to conduct sample collection among the Vedda clans. We also sincerely thank Mr Eranda Dissanayake of the Dambana Information Centre, Central Cultural Fund, for coordinating the sample collection process. This work received partial financial support from the National Research Council (NRC) through Grant no: 16-023, with additional support from Genetech, Colombo.

## Ethical Declaration

Ethical approval for this research was obtained from the Ethics Review Committee of the Institute of Biology, Sri Lanka (Reference ERC IOBSL 269 08 2022).

